# INSULIN GROWTH FACTOR I AND ITS RECEPTOR ARE ANTAGONISTIC MODULATORS OF GLUCOSE HANDLING BY ASTROCYTES

**DOI:** 10.1101/023556

**Authors:** E. Hernandez-Garzón, A.M. Fernandez, A. Perez-Alvarez, S. Mederos, P. Perez-Domper, P. Bascuñana, R.F. de la Rosa, M. Delgado, M.A. Pozo, A. Miranda-Vizuete, D. Guerrero-Gomez, E. Moreno, P.J. McCormick, A. Santi, L. Genis, A. Trueba, C. Garcia-Caceres, M.H. Tschöp, A. Araque, G. Perea, E.D. Martin, I. Torres Aleman

## Abstract

Reducing insulin-like growth factor I receptor (IGF-IR) levels or administration of IGF-I show beneficial effects in the brain. We now provide evidence to help resolve this paradox. The unliganded IGF-IR inhibits glucose uptake by astrocytes while its stimulation with IGF-I, in concert with insulin activation of the insulin receptor, produces the opposite effect. In vivo imaging showed that shRNA interference of brain IGF-IR increased glucose uptake by astrocytes while pharmacological blockade of IGF-IR reduced it. Brain ^18^FGlucose-PET of IGF-IR shRNA injected mice confirmed an inhibitory role of unliganded IGF-IR on glucose uptake, whereas glucose-dependent recovery of neuronal activity in brain slices was blunted by pharmacological blockade of IGF-IR. Mechanistically, we found that the unliganded IGF-IR retains glucose transporter 1 (GLUT1), the main glucose transporter in astrocytes, inside the cell while IGF-I, in cooperation with insulin, synergistically stimulates MAPK/PKD to promote association of IGF-IR with GLUT 1 via Rac1/GIPC1 and increases GLUT1 availability at the cell membrane. These findings identify IGF-I and its receptor as antagonistic modulators of brain glucose uptake.

## Introduction

IGF-I is generally considered a neuroprotective and pro-cognitive factor and has even been proposed as therapy for Alzheimer’s dementia and other neurodegenerative diseases [1] However, reduced IGF-I receptor levels have been shown to ameliorate pathology in animal models of Alzheimer’s disease [2]. This poses the paradox that either increasing IGF-I or reducing its receptor leads to beneficial actions in the brain [1, 3]. While these apparently contradictory observations remain largely unexplained (but see [4]), one possibility is that ligand-independent actions of IGF-IR antagonize the actions of IGF-I, as recently reported for apoptotic signaling through insulin receptor (IR) and IGF-IR [5]. Other possibilities include a network of complex interactions among insulin-like peptides (ILPs) and their receptors, as described in the nematode *Caenorhabditis elegans* with a single insulin-like receptor and multiple ILP ligands that may show even opposite activities [6].

In our search for an explanation of these discrepancies we analyzed the role of insulin and IGF-I receptors and their ligands on glucose handling by the brain. ILPs are important modulators of cell energy balance, a key aspect in tissue homeostasis that could in theory form part of neuroprotection by ILPs and remains little explored. Yet, while there is evidence that IGF-I affects glucose metabolism in the brain [7-9], the role of insulin is not clear [10], even though the brain widely expresses IR [11]. Only under pathological circumstances the gluco-regulatory actions of insulin on the brain manifest [12]. Moreover, both endothelial cells and astrocytes, the main cellular constituents of the blood-brain barrier (BBB) express glucose transporter 1 (GLUT1) as their main facilitative transporter [13], and GLUT1 is considered largely insulin insensitive [14].

We now describe that the unliganded IGF-I receptor reduces glucose uptake by the brain by inhibiting glucose capture by astrocytes. The intrinsic inhibitory action of IGF-IR is countered by the concerted action of insulin and IGF-I. Opposite ligand-dependent vs ligand-independent roles of IGF-IR on brain glucose uptake may help understand the reported paradoxical actions of this growth factor system in the brain [3].

## Results

### Opposing roles of insulin-like growth factor I receptor and insulin/IGF-I on glucose uptake

We determined whether insulin/IGF-I and their receptors display opposite effects on brain glucose uptake. As described over a century ago [15], neuronal activity increases blood flow through neuro-vascular coupling at the BBB and in this way blood nutrients, including glucose, are taken up by the active brain region. Because endothelial cells and astrocytes are the key cell types at the BBB involved in blood glucose uptake, we first examined whether insulin and IGF-I modulate glucose capture by these two type of cells using the fluorescent glucose analog 6-NBDG [16]. Whereas neither insulin nor IGF-I modulated glucose uptake in these cells, the combined addition of insulin and IGF-I (I+I) stimulated it in astrocytes, -but not in endothelial cells (Fig 1A and S1A). Glucose uptake by neurons, the major energy consumers of the brain, was not affected by insulin peptides under these experimental conditions (S1B Fig).

**Figure 1:**
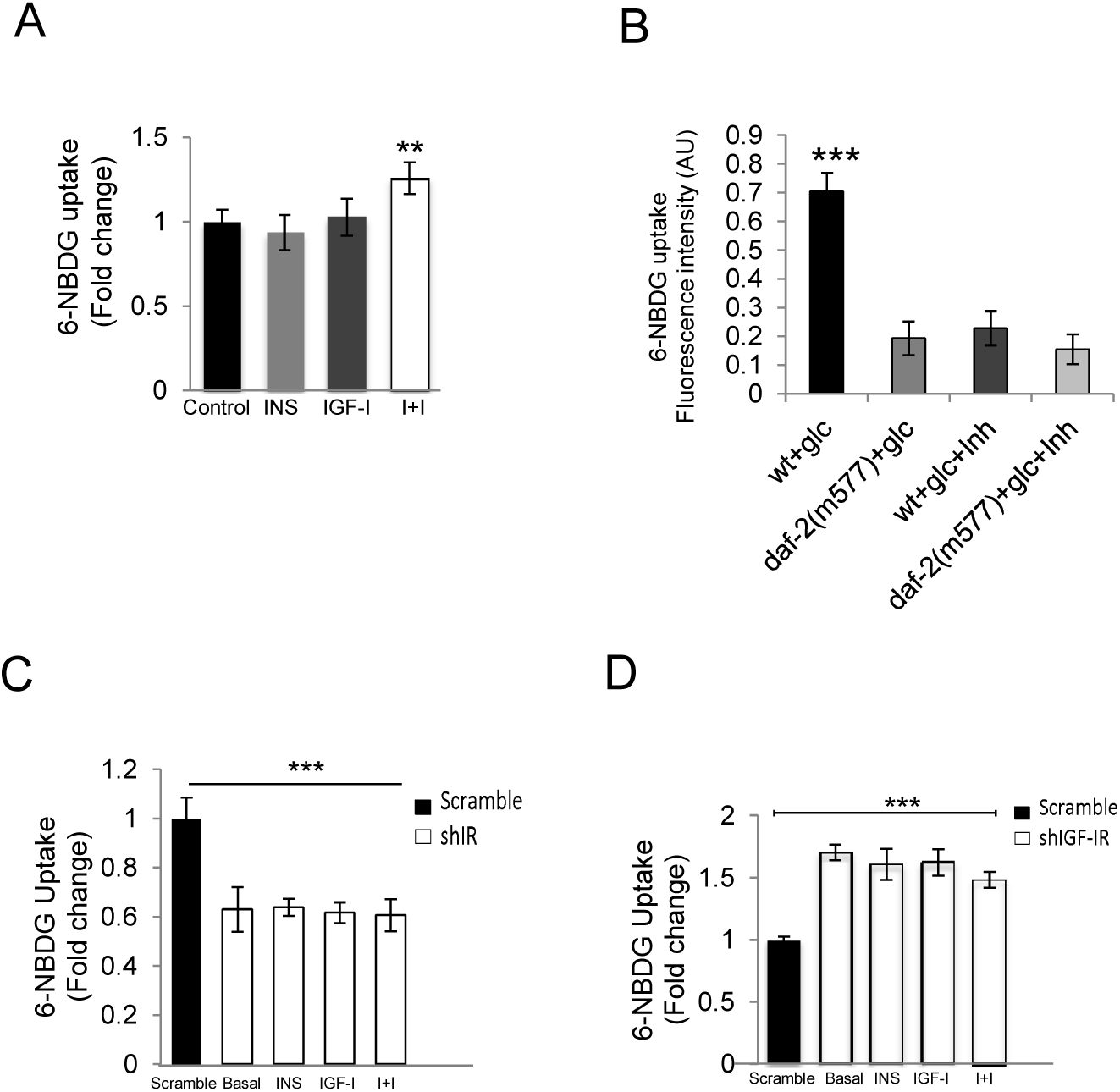
Modulation by ILPs and their receptors of glucose uptake. **A**, Astrocytes exposed to insulin (INS) plus IGF-I (I+I) showed increased accumulation of the fluorescent glucose analog 6-NBDG (n=10; **p<0.01 vs basal control levels). **B**, Glucose uptake in *C. elegans* is regulated by the insulin-like receptor DAF-2. Glucose (glc) uptake is reduced in daf-2 (m577) and daf-2 (e-1370, not shown) mutants and in wild type (wt) worms administered with the DAF-2 inhibitor NT219 (inh). Fluorescence values obtained for the 6-NBDG treated animals were corrected by the mean fluorescence of non-treated controls of the same genetic background. ***p<0.001 vs all other groups, (n=35 per group). **C**, Depletion of IR by shRNA interference elicits the opposite effect: a decrease in basal uptake of 6-NBDG and abrogation of the increase elicited by I+I (n=6; ***p<0.001 vs scramble-transfected astrocytes). **D**, Depletion of IGF-IR in astrocytes by shRNA interference increases basal uptake of 6-NBDG and no further increases are seen after I+I (n=6; ***p<0.001 vs scramble-transfected astrocytes).

We next determined the role of IGF-I and insulin receptors on glucose uptake. Since neuroprotective effects of reduced ILP receptor activity were first described in *C elegans* [17], we assessed whether DAF-2, the single nematode insulin-like receptor is involved in glucose handling as the receptors of the insulin family display highly conserved activities. To our knowledge, whether DAF-2 plays a role in glucose homeostasis in the worm has not yet been explored, although recent observations indirectly favor its involvement [18]. In two *daf-2* hypomorphic alleles, *daf-2* (*e1370*, data not shown) and *daf-2 (m577)*, representative of each of the two phenotypic classes of *daf-2* alleles [19], whole body 6-NBDG uptake was significantly lower than in wild type controls (Fig 1B), suggesting that DAF-2 behaves as the mammalian insulin receptor [20] regarding body glucose handling. Accordingly, inhibition of *daf-2* by pharmacological blockade mimicked the phenotype of *daf-2* mutants in wild type worms (Fig 1B), similarly to previous observations in longevity traits [21].

We then analyzed the role of the two mammalian tyrosine-kinase insulin receptors; i.e.: IGF-IR and IR in glucose uptake by reducing their levels using shRNA interference in astrocytes (S1C,D Fig). We chose these cells as they were the only ones that responded to insulin and IGF-I and express both types of receptors (S2A Fig). We found that reduction of either one modified glucose uptake by astrocytes, but in opposite directions. Similarly to daf2, decreased IR significantly decreased glucose uptake (Fig 1C), whereas reduction of IGF-IR resulted in increased glucose uptake (Fig 1D). Opposing actions of IR and IGF-IR on glucose uptake were confirmed by the observation that when both receptors were reduced the effects cancelled each other (S2B Fig). Moreover, reduction of either IR or IGF-IR was sufficient to block the combined action of I+I (Fig 1C,D).

### Cooperation between insulin and IGF-I require insulin and IGF-I receptors

The fact that insulin requires IGF-I to stimulate glucose uptake, and only in astrocytes, may help explain the difficulty to establish a gluco-regulatory action of this hormone in the brain. Therefore, we explored possible mechanisms involved. As glucose uptake was stimulated only when insulin and IGF-I were added simultaneously and the effect disappeared when either receptor was knocked down (Fig 1C,D), an interaction of both hormones through their respective receptors was deemed probable. Using cultured astrocytes we inhibited either IR or IGF-IR with the specific antagonists S961 and PPP, respectively. In both cases the effects of I+I were blocked (Fig 2 A,B). Expression of dominant negative forms of either IGF-IR or IR similarly abolished the stimulatory actions of I+I (S2C,D Fig). Therefore, both receptors need to be activated to stimulate astrocyte glucose uptake. To determine whether this in vitro observations bear a functional impact we took advantage that neurons release insulin and IGF-I while depolarized [22, 23], and that during high energy demands neurons rely on astrocytes for additional energy supply [24]. Thus, we recorded excitatory synaptic transmission (fEPSPs) in cortical slices under changing glucose concentrations and, as expected, hypoglycemia gradually reduced fEPSPs (Fig 2C). Upon glucose re-addition to the medium, fEPSPs almost fully recovered under control conditions while in the presence of drugs blocking insulin or IGF-I receptors, recovery was significantly reduced (Fig 2C). These observations provide indirect support of findings in cultured astrocytes and favor the notion that both receptors need to be active for astrocytes to properly supply energy to neurons.

**Figure 2:**
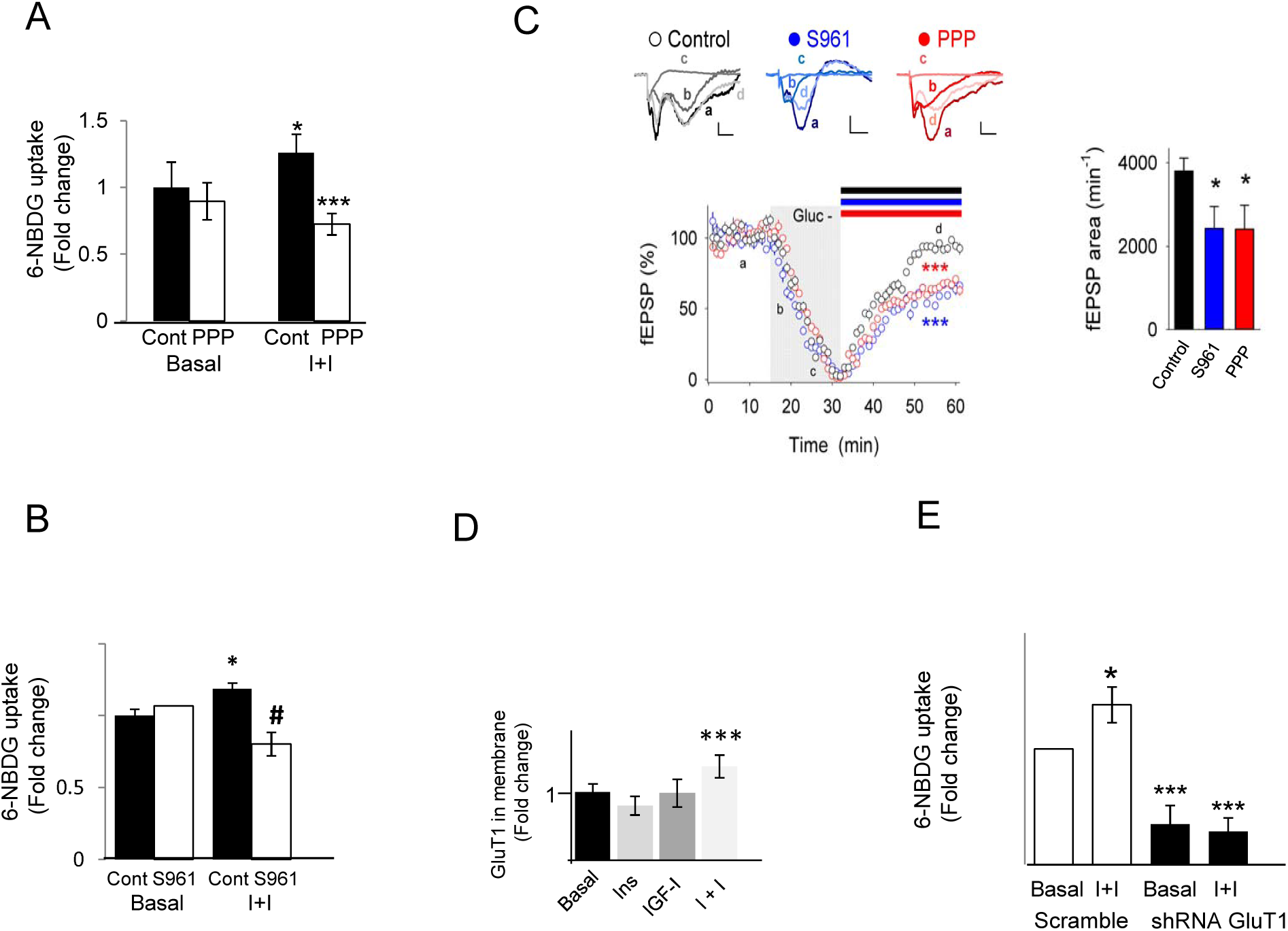
Ligand-dependent stimulatory actions of IGF-IR and IR on glucose uptake by astrocytes. **A, B**, Stimulated uptake of 6-NBDG after I+I is abrogated by specific IGF-IR antagonist (PPP), or IR antagonist (S961; n=6; *p<0.05 vs basal control, **p<0.01 vs I+I alone and #p<0.05 vs I+I alone). **C**, Insulin and IGF1 receptor activation is involved in recovery of hypoglycemic-induced depression of synaptic transmission. *Left top*, Representative fEPSP traces in response to glucose deprivation in control conditions (black), in the presence of the insulin receptor inhibitor S961 (blue), or the IGF-I receptor inhibitor PPP (red). a-d denote synaptic responses before glucose deprivation (baseline, a), during hypoglycemia onset (b), hypoglycemia plateau (c), and recovery after adding glucose-ACSF (d) in control (black bar), S961, or PPP (blue and red bars, respectively) conditions. *Left bottom*, Time course of fEPSP slope changes during hypoglycemia (gray bar) in control (white; n=10 slices), PPP (10 □M; red; n=10) or S961 (10 nM; blue; n=12) conditions. *Right*, Relative changes of the area for the fEPSP slope recovery after addition of glucose-ACSF under control (black), PPP (red) or S961 (blue) conditions. Data are presented as the mean values ± SEM. * p<0.05 and ***p<0.001 vs control. **D**, The amount of GLUT1in the cell membrane is significantly increased by the combined addition of I+I in astrocytes, but not by either of them alone, as determined by flow cytometry of GFP-tagged GLUT1 (n=6; ***p<0.001 vs control). **E**, Reduction of GLUT1 by shRNA interference drastically reduces 6-NBDG accumulation by astrocytes and abrogates the stimulatory effects of I+I (n=4; *p<0.05 and ***p<0.001 vs basal control levels).

As blood-borne glucose is captured by the brain through GLUT1, the most abundant type of facilitative transporter in brain endothelia and astrocytes [25], we determined whether astrocytic GLUT1 is involved in glucose uptake after I+I. Indeed, the amount of GLUT1 translocated to the astrocyte cell membrane was significantly increased in the presence of I+I (Fig 2D and S3A). Furthermore, when GLUT1 levels in astrocytes were reduced with shRNA (S3B Fig), basal glucose uptake was drastically reduced and the stimulatory action of I+I was no longer seen (Fig 2E).

Intracellular signaling pathways downstream of I+I were then examined. We measured phosphorylation of AKT and MAPK because these two kinases form part of the canonical signaling of both insulin and IGF-I receptors. As shown in Fig 3A, phospho-MAPK, but not phospho-AKT (S3D Fig) was synergistically elevated in astrocytes by I+I. As potential downstream mechanisms we focused on PKD, an atypical PKC isoform that may act downstream of MAPK to regulate translocation of glucose transporters [26], and associates to IGF-IR but not to IR [27] (see below). PKD was synergistically stimulated by I+I, as evidenced by increased levels of phosphoSer-PKD (Fig 3B), in a MAPK-dependent manner (Fig 3C). Further, the PKD inhibitor CID755673 abrogated increased glucose uptake by astrocytes in response to I+I (Fig 3D). Collectively, these data support a cooperative action of insulin and IGF-I in glucose handling by astrocytes through synergistic activation of MAPK_42-44_ and PKD.

**Figure 3:**
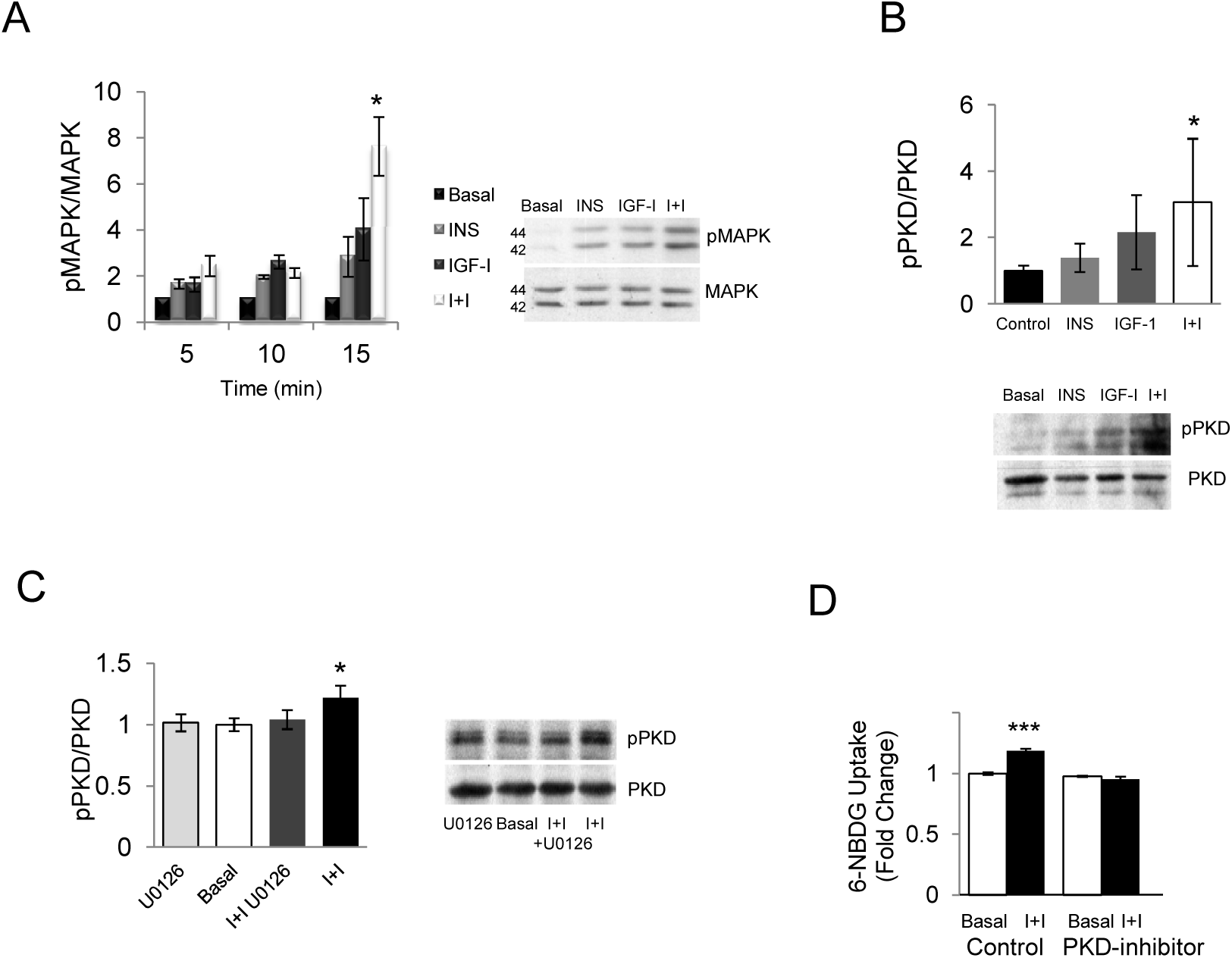
**A**, Levels of phosphorylated MAPK_42-44_ (pMAPK) were synergistically elevated by co-addition of insulin and IGF-I to astrocytes. Levels of pMAPK were normalized by total levels of MAPK_42-44_. Representative blot after 15 minutes of stimulation with insulin, IGF-I or both is shown. Bars show quantification of the ratio pMAPK/MAPK after 5, 10, and 15 minutes of stimulation (*p< 0.05 vs insulin or IGF-I alone, n=3). **B**, PKD phosphorylation is synergistically increased by I+I. Representative blot after 15 minutes of stimulation with insulin, IGF-I or both is shown. Bars show quantification of the ratio pPKD/PKD (*p<0.05 vs basal; n=3). **C**, Increased PKD phosphorylation after I+I depends on MAPK as in the presence of the MAPK inhibitor U0126 the increase was abolished. Representative blot after 15 minutes of stimulation with I+I, U0126 or I+I+U0126 is shown. Bars show quantification of the ratio pPKD/PKD (*p<0.05 vs basal; n=3). **D**, Stimulated uptake of 6-NBDG after I+I is abrogated by the PKD inhibitor CID (10 μM) (*p<0.05 vs basal control; n=6).

### IR and IGF-IR have opposite actions on glucose uptake

Modulation of basal glucose uptake in astrocytes after reduction of either IR or IGF-IR probably reflects ligand-independent actions of these receptors, as previously reported [5]. Indeed, decreasing IR levels by shRNA interference reduced GLUT1 mRNA (Fig 4A) and decreased GLUT1 in the cell membrane (Fig 4B). In agreement with these observations we found reduced levels of brain GLUT1 mRNA in mice lacking IR in astrocytes (Fig 4C). Hence, IR increases GLUT1 expression, and consequently GLUT1 protein levels at the cell membrane, in a ligand-independent manner. On the other hand, reduction of IGF-IR by shRNA did not affect GLUT1 mRNA levels (Fig 4A), but resulted in increased amounts of GLUT1 at the cell membrane of astrocytes (Fig 4B). The latter observation suggests that the IGF-I receptor exerts a ligand-independent inhibitory effect on glucose uptake by reducing the availability of GLUT1 at the cell membrane.

**Figure 4:**
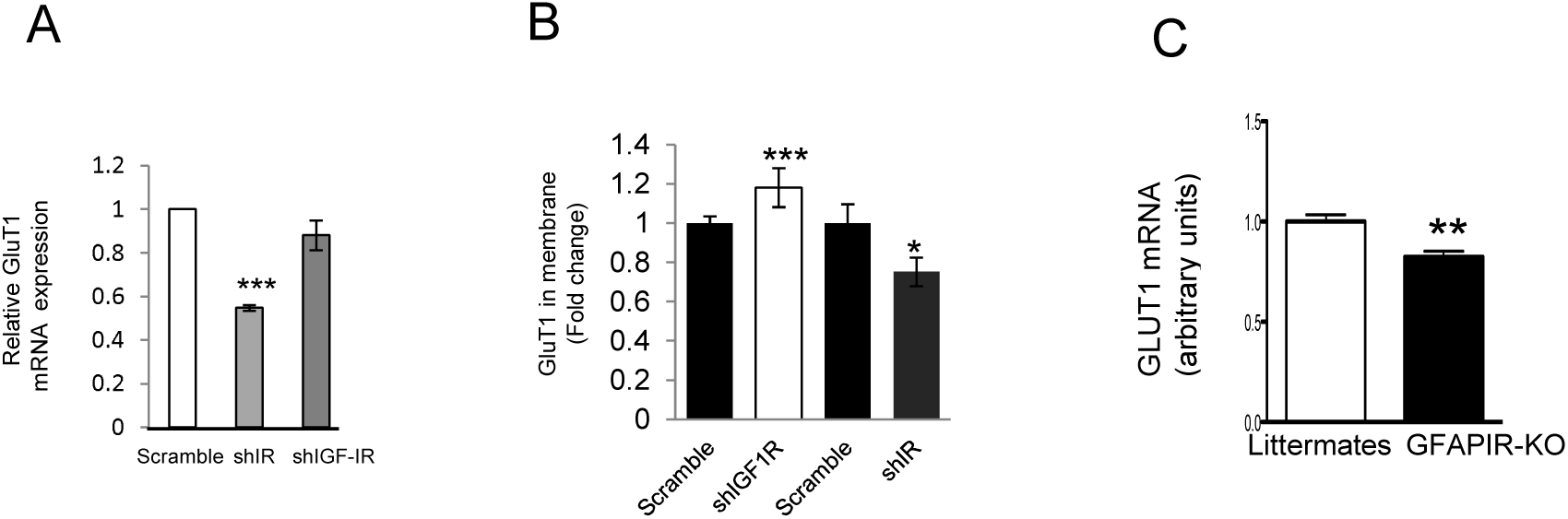
Opposite regulation of glucose uptake by IGF-IR and IR. **A**, Depletion of IR, but not IGF-IR by shRNA interference decreases the amount of GLUT1 mRNA as determined by qPCR (n=3; ***p<0.001 vs scramble-transfected astrocytes). **B**, Reduction on of IGF-IR increased the amount of GLUT1 in the cell membrane whereas depletion of IR decreased it (n=4; ***p<0.001, and *p<0.05 vs respective scramble). **C**, Brain levels of GLUT1 mRNA are significantly reduced in mice lacking insulin receptors in astrocytes (IR-KO As) as compared to control littermates (**p<0.01 vs littermates; n=4).

### Unliganded and liganded IGF-IR has opposite effect on glucose uptake

The above in vitro observations indicate that the unliganded IGF-IR antagonizes glucose uptake whereas IGF-IR is activated by IGF-I to stimulate glucose uptake in concert with insulin. Hence, we conducted further experiments to determine whether this dual, antagonistic action is also observed in vivo. We took advantage of the fact that upon somatosensory stimulation brain glucose uptake is predominantly mediated by astrocytes [28]. Using ^18^F fluoro-2 deoxy-glucose PET to measure brain glucose handling we injected mice with a lentiviral vector expressing IGF-IR shRNA into one side of the somatosensory cortex whereas the contralateral side was injected with the same viral vector expressing a scrambled shRNA (S4A Fig). Upon bilateral stimulation of the whiskers -that increases glucose flux into the somatosensory cortex, significantly greater levels of glucose uptake were found in the side injected with IGF-IR shRNA after normalizing to the increased uptake of the contralateral scramble-injected site (Fig 5A and S4B). Despite the limitations of PET analysis in mice [29], these results support the observation that reducing brain IGF-IR levels enhances glucose uptake by astrocytes. To confirm that astrocytes were the cells involved in glucose accumulation we used in vivo fluorescence microscopy (S5 Fig). After reduction of IGF-IR in the mouse somatosensory cortex by viral IGF-IR shRNA expression as before, larger increases in glucose uptake (detected with 6-NBDG) in somatosensory cortex astrocytes (identified with the astrocyte-specific fluorescent red marker SR101) were elicited upon stimulation of the whiskers as compared to scramble RNA-injected mice (Fig 5B,C). No changes were seen after intracortical injection of IR shRNA (not shown). This reinforces a ligand-independent inhibitory role of IGF-IR on glucose uptake by astrocytes.

**Figure 5:**
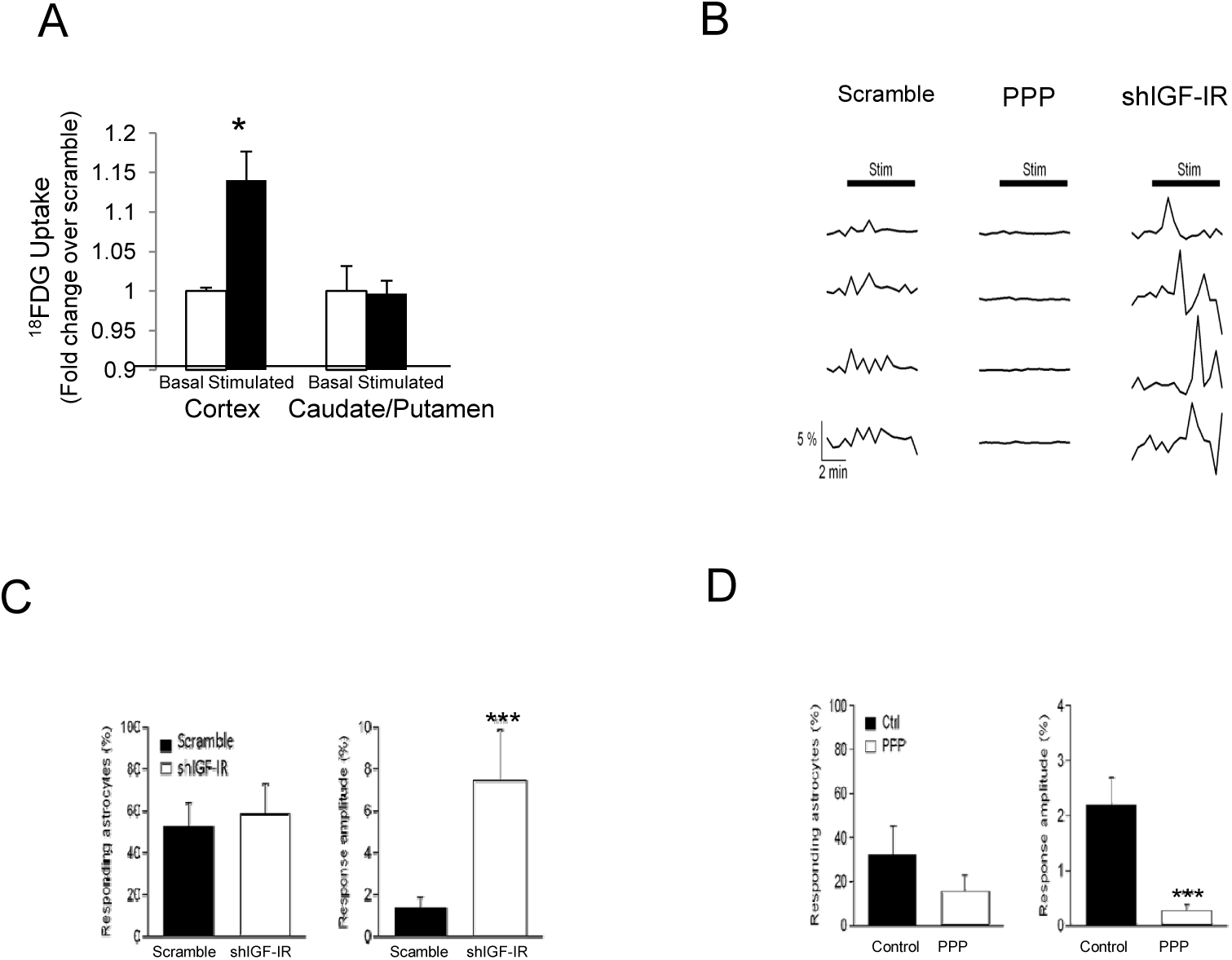
Opposite regulation of glucose uptake by the unliganded and liganded IGF-IR. **A**, Adult mice were unilaterally injected with lentival particles expressing shRNA IGF-IR (n=5) in one side of the somatosensory cortex and with scramble shRNA-expressing viral particles in the contralateral side (see Suppl Fig 4A). After allowing 2 weeks of recovery animals were submitted to PET scans and after basal measurements of ^18^F-FDG uptake they were stimulated in their whiskers. Mice injected with shRNA IGF-IR showed significantly enhanced uptake in the somatosensory cortex in response to bilateral whisker stimulation as compared to the scramble injected side. A reference unstimulated area, the caudate putamen, was analyzed to determine region specificity in increased glucose uptake (*p<0.05 vs basal). **B**, Representative 6-NBDG fluorescence traces of astrocytes of somatosensory cortex under basal conditions, after bathing the cortical surface with the IGF-I receptor blocker PPP (middle), or after injection of IGF-IR shRNA in the somatosensory cortex. **C**, Mice injected with IGF-IR shRNA showed no changes in the number of responding astrocytes (left histograms) but displayed a great increase in glucose uptake (measured as amplitude of the fluorescence response of 6-NBDG; right histograms, n= 139; p<0.001 vs scramble shRNA; n= 123). **D**, Cortical dministration of PPP (n=188) did not modify the number of astrocytes responding to sensory stimulation (left histograms) while the uptake of glucose was markedly diminished (right histograms; p<0.0001 vs control; n= 108).

We next analyzed whether the ligand-dependent actions of IGF-IR could also be evidenced using this in vivo approach. As pharmacological inhibition of IGF-IR with PPP blocked glucose uptake in vitro (Figure 2A), we locally infused this receptor antagonist into the somatosensory cortex of wild type mice. In PPP-infused mice, no increases in glucose uptake by astrocytes were seen in response to whisker stimulation (Fig 5B,D), a procedure that activates both IGF-I and insulin receptors in somatosensory cortex (see S6 Fig). The latter agrees with reduced neuronal activity after PPP infusion in somatosensory cortex during whisker stimulation [30].

### Mechanisms of cooperation between insulin and IGF-I

Since GLUT1 is involved in glucose uptake promoted by I+I, we examined possible underlying mechanisms. IGF-IR associates with GIPC (GAIP-interacting protein, C terminus) [31, 32], a scaffolding protein that participates in protein trafficking and binds to many partners, including GLUT1 [32]. We hypothesized that GIPC may simultaneously interact with IGF-IR and GLUT1 through its PDZ domain because GIPC can dimerize [33]. In support of this possibility we found that IGF-IR co-immunoprecipitates not only with GIPC, as already reported, but also with GLUT1, whereas GIPC, as expected, co-immunoprecipitates also with GLUT1 (Fig 6A). Proximity ligation assays (PLA, Fig 6B) and confocal analysis (not shown) confirmed an interaction between IGF-IR and GLUT1 in astrocytes.

**Figure 6:**
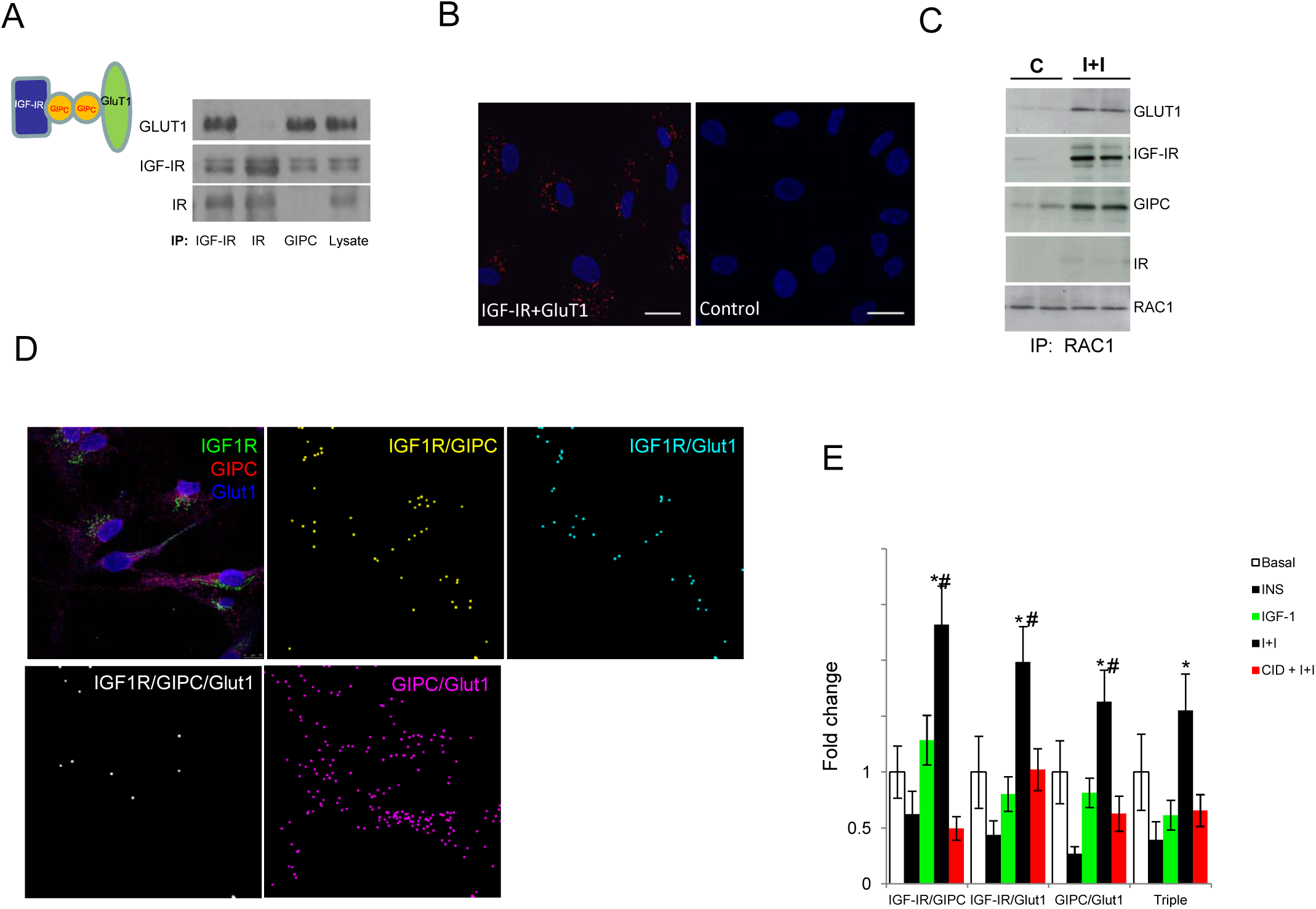
Mechanisms of cooperativity of insulin and IGF-I to stimulate glucose uptake. **A**, Analysis of interactions of IGF-IR with GLUT1 and GIPC using reciprocal immunoprecipitations and sequential blotting show that the 3 proteins interact with each other whereas only IGF-IR interacts with IR. Drawing: proposed interaction of GLUT1 with IGF-IR through GIPC. **B**, Proximity ligation assays (PLA) in astrocyte cultures show an interaction of IGF-IR with GLUT1 (red dots in left micrograph) that is absent when the IGF-IR antibody is omitted (Control). DAPI staining of astrocyte nuclei in blue. Bars are 20μm. **C**, Addition of I+I reinforces the association of IGF-IR with GIPC and GLUT1 and promotes the association of Rac1 to the complex. IR does not associate to Rac1 either. Co-immunoprecipitation was performed with anti-Rac1. **D**, Representative confocal microscopy images showing staining for IGF-IR, GIPC1, and GLUT1 in cultured astrocytes. Double and triple staining particles were scored. Representative images of double- and triple-particle staining are codified with different colors and shown 4X magnified to better illustrate them. Bar is 10 μm. **E**, Co-addition of insulin and IGF-I significantly stimulated the relative amount of IGF-IR/GIPC, IGF-IR/ GLUT1, GIPC/GLUT1, and IGF-IR/GLUT1/GIPC-immunopositive particles as compared to insulin or IGF-I given alone. When PKD is inhibited with CID, the pattern of immunopositive particles after I+I drastically changes. The different types of particles were scored using image J analysis software. N= 22 cells for control, 28 for insulin, 33 for IGF-I, 20 for I+I, and 46 for I+I+CID (*p<0.05 vs insulin, IGF-I and I+I+CID, and # p<0.05 vs basal).

We then examined whether the interactions between IGF-IR, GLUT1 and GIPC1 are modified by addition of insulin + IGF-I. Confocal analysis (Figure 6D) showed a specific pattern of changes in the number of double and triple-containing puncta after I+I stimulation, as compared to basal conditions or after insulin or IGF-I alone (Figure 6E). Addition of insulin greatly decreased all types of immunostained puncta, IGF-I slightly modified some of them, whereas I+I synergistically increased the interactions IGF-IR/GLUT1, IGF-IR/GPIC and GLUT1/GIPC (Fig 6E, p<0.05 vs basal conditions). PLA analysis of GLUT1/IGF-IR interactions after I+I confirmed an increase of: 2.54 ± 0.59-fold over control (p<0.05; n=4). To link the pattern of puncta mobilization after I+I with glucose uptake we inhibited the latter by inhibiting PKD with CID (Fig 3D). CID drastically altered the pattern of puncta mobilization elicited by I+I (Fig 7C), supporting a role of this pattern in glucose uptake. Further confirmation of a role of the observed reinforced protein-protein interactions in glucose uptake after I+I was carried out with co-immunoprecipitation assays. An increased interaction of IGF-IR with GLUT1 and also with GIPC was found after I+I (S7A Fig). Because GIPC modulates the activity of the small GTPase Rac1 [34], that is involved in translocation of glucose transporters from intracellular compartments to the cell membrane [35] we also analyzed its possible involvement in the action of insulin and IGF-I. We found that I+I promoted the interaction of Rac 1 with GIPC, GLUT1 and IGF-IR (Figure 6C and S7A), an interaction not seen under basal conditions. Neither GLUT1, Rac1, or as previously shown [36], nor GIPC co-immunoprecipitate with the insulin receptor (Fig 6A,C).

**Figure 7:**
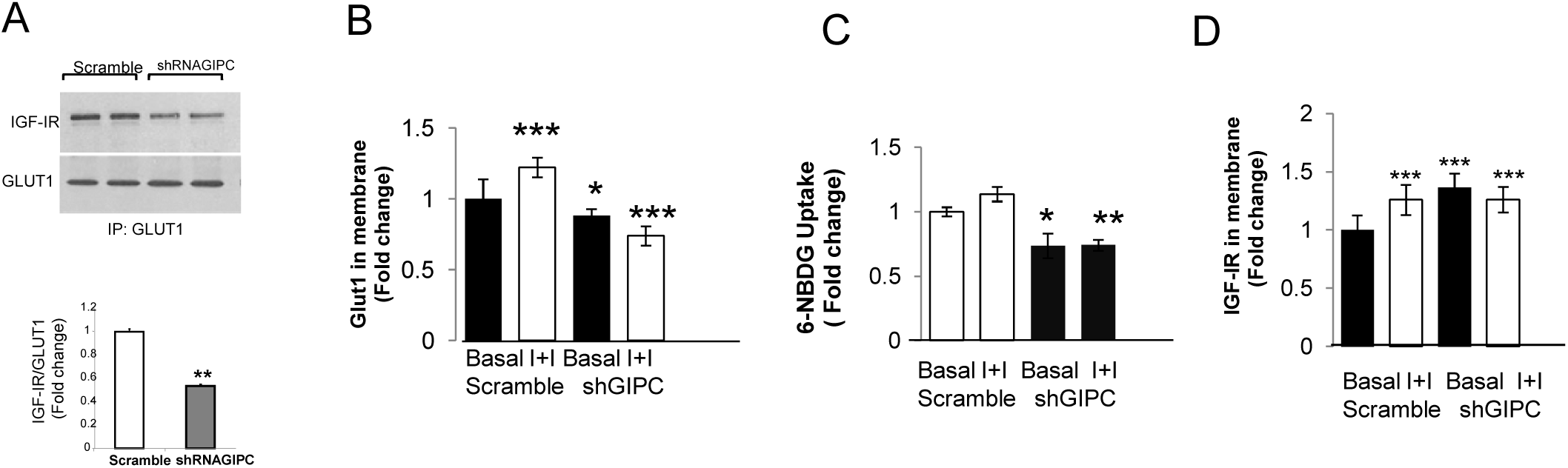
IGF-IR interacts with GLUT1 through GIPC1. **A**, Reduction of GIPC1 with shRNA interference resulted in significantly less IGF-IR bound to GLUT1. Representative blot and quantitation bars are shown (n=4; **p<0.01). **B**, Reduction of GIPC1 results in reduced levels of GLUT1 in the membrane and abrogation of the stimulatory effects of I+I (n=6; *p<0.05 and ***p<0.001 vs basal control levels). **C**, Reduced GIPC1 also impedes stimulation by I+I of 6-NBDG uptake by astrocytes (n=6; *p<0.05 and **p<0.01 vs control levels). **D**, Conversely, reduction of GIPC1 increases the amount of IGF-IR in the cell membrane under basal conditions without any further increase after I+I (n=6; ***p<0.001 vs control levels).

To further establish a role of GIPC in the interaction between IGF-IR and GLUT1 we reduced GIPC levels in astrocytes using GIPC shRNA (S7B Fig) and found that the interaction between IGF-IR and GLUT1 was significantly reduced (Fig 7A). GIPC is required to sort GLUT1 to the plasma membrane [37]. Accordingly, GIPC reduction by RNA interference decreased the amount of GLUT1 in the cell membrane (Fig 7B) and, as a result, basal glucose uptake was reduced and stimulation by I+I impaired (Fig 7C). Conversely, the amount of IGF-I receptor at the cell membrane measured by flow cytometry was increased by GIPC shRNA and was not further increased by I+I (Fig 7D). Opposite changes in IGF-IR and GLUT1 levels at the cell membrane when GIPC is reduced supports the notion that GIPC participates in regulating their membrane traffic. Indeed, GIPC is involved in GLUT1 sorting to the cell membrane [37] and in IGF-IR recycling [38]. This agrees with the observation that enhanced glucose uptake after I+I recruits to the cell membrane not only GLUT1, as can be expected for a glucose transporter, but also IGF-IR. To reconcile the presence of higher levels of IGF-IR in the cell membrane after I+I with an involvement of GIPC, whose depletion increases IGF-IR levels at the cell membrane (Fig 7D), we hypothesized that the interactions of GIPC with IGF-IR and GLUT1 are directed by I+I in a way that promotes GLUT1 translocation to the cell membrane and at the same time IGF-IR recycling (S7C Figure).

Collectively, these observations are compatible with the notion that the concerted activation of insulin and IGF-I receptors increases glucose uptake in astrocytes through a synergistic activation of PKD that in turn stimulates IGF-IR recycling and GLUT1 translocation to the membrane by modulating the interaction of IGF-IR with GLUT1 through GIPC and Rac 1.

## Discussion

The present observations indicate that IGF-IR has an intrinsic inhibitory action on astrocytic glucose uptake opposite to an intrinsic stimulatory activity of IR. Therefore, glucose uptake in astrocytes may in part be determined by a balance between IGF-IR and IR levels. This suggests that physiological and pathological processes impacting on brain levels of insulin and IGF-I receptors will influence glucose handling by the brain in opposite directions. In addition, insulin and IGF-I bind to IR and IGF-IR to stimulate glucose uptake. In this way, insulin and IGF-I counter intrinsic IGF-IR activity and potentiate intrinsic IR actions. These complex set of interactions may help understand paradoxical effects of ILPs and their receptors in neuroprotection. For example, in terms of functional impact it would not be the same to reduce IGF-IR activity than to reduce IGF-I activity. In other words, either reduction of IGF-IR or increasing insulin peptides will similarly enhance glucose uptake by the brain, which fits well with previously observed beneficial actions of decreased IGF-IR [2] or increased IGF-I [39] on brain function. While in primitive organisms such as C *elegans* a single ILP receptor is modulated by many different ligands, even in an antagonistic fashion [40], the acquisition of new ILP receptors in vertebrates has allowed the appearance of novel interactions among them. In our view, reported actions of ILPs in invertebrates should not be immediately inferred to be similar in vertebrates. The corollary of these observations is that invertebrate models of ILP physiology in mammals should take into account the existence of two tyrosine kinase ILP receptors, IGF-IR and IR, not present in invertebrates that may display cooperative [5] or opposing activities (present observations), depending on biological context.

A possible gluco-regulatory role of insulin in the brain has remained elusive probably because as our observations indicate, glucose uptake in astrocytes is regulated by this hormone in cooperation with IGF-I. Insulin and IGF-I recruit their respective receptors and synergistically stimulate a MAPK/PKD pathway. In turn, this pathway promotes the interaction of IGF-IR with GLUT1 and GIPC and the association of the GTPase Rac1 to translocate GLUT1 to the cell membrane. Thus, glucose uptake is enhanced by the concerted action of insulin and IGF-I because GLUT1 availability at the astrocyte cell membrane is increased. Significantly, in cortical slices that produce both IGF-I and insulin (not shown), inhibition of either receptor was sufficient to abrogate recovery of neuronal activity after hypoglycemia. In line with this, recent in vitro data confirm a role of insulin and IGF-I in astrocytic glucose metabolism [41]. Altogether these set of observations support a role of an interactive network of insulin peptides and their receptors in modulating glucose handling by astrocytes, adding further complexity to the role of astrocytes in brain energy economy [42].

Because glucose uptake by the brain deteriorates during aging and its associated pathologies [43], these observations potentially open also a new path for a future therapeutic rationale for healthy as well as pathological aging. Indeed, our observations provide a direct link between insulin and IGF-I resistance in Alzheimer’s (AD) brains [44] and the well established pattern of glucose dysregulation present as a characteristic alteration of AD [45]. Future work should examine major components of the insulin/IGF-I dependent pathway described herein for brain glucose uptake in experimental models of normal and pathological aging, as illustrated by the recently described role of endothelial GLUT1 in AD [46].

## Materials and Methods

**Animals**: Adult and new-born C57BL6/J mice and mutant hGFAP-CreERT2 mice were used.

**Cell cultures and transfections**: Astrocytes from postnatal 3-days forebrain (>95% GFAP^+^ cells), granule neurons from P7 cerebella (>99% β3-tubulin^+^ cells), and brain endothelial cells (P7-10 brain, >90% CD31^+^ cells) were used. Astrocytes were electroporated using a Nucleofector Kit (Amaxa). Transfection efficiency was 60–80%. For viral transduction a three-plasmid lentiviral system was used.

**Glucose Uptake :** Glucose uptake was determined with the fluorescent glucose analog 6-NBDG by flow cytometry. **Protein Translocation Assay:** Translocation of GLUT1 and IGF-IR to the cell membrane was evaluated also by flow cytometry using anti IGF-IR or anti Flag M2 antibodies. GLUT1-IGF-IR interactions were detected with Duolink II in situ PLA detection (OLink). For confocal analysis, pictures were taken with a 63X oil immersion objective. For each field, a series of 8 to 12 Z stacks with a step size of 0.29 μm in each channel were acquired. Number of immunopositive particles was determined taking 20-34 cells/5 fields.

**qPCR:** For quantitative PCR, cDNA was amplified using TaqMan probes for GLUT1, GluT4, IGF-IR or IR, and 18S as reference control.

***In vivo* experiments**: Lentiviral particles (2 μl/mouse) expressing shRNA against IGF-I receptor receptor were administered at the stereotaxical coordinates: -1.06mm from bregma and -1mm lateral. For in vivo glucose uptake mice were anesthetized, astrocytes labelled with sulforhodamine 101 (injected ip), and through their femoral artery animals received 6-NBDG. A 4 mm craniotomy around the somatosensory cortex was made. The IGF-IR blocker PPP or the vehicle was added to the exposed somatosensory cortex. Whiskers were stimulated with 100 ms puffs of air and tails pinched at 2 Hz with forceps. Imaging was acquired with a confocal laser scanning microscope. Images were aligned over previous ones with Align Slice (Image J). Fluorescence intensity was measured in a ROI limited to the somatic area and expressed as relative fluorescence changes (ΔF/F0), where F0 is the mean of the baseline period.

**^18^F-FDG PET:** ^18^F-FDG PET was used to measure brain glucose handling following standardized procedures. PET acquisition was 20 min, followed by computed tomography. After ^18^F-FDG uptake was calculated, the activity of each left hemisphere region was normalized to its homologous region in the right hemisphere and expressed as proportional uptake (left/right).

**Slice recordings:** Cortical slices from wild type mice were stimulated with bipolar electrodes placed on layer 4 of somatosensory cortex and recording glass microelectrodes in layer 2. A stimulus intensity which evoked half-maximum amplitude fEPSPs was used. Baseline responses were recorded and test stimuli given at 0.1 Hz.

**6-NBDG incorporation in *Caenorhabditis elegans:*** Analysis of glucose uptake in wild type and daf-2 mutant *C. elegans* was carried out using 6-NBDG. Worms were allowed to stain for 120 min at room temperature in the dark with gently shaking. Animals were mounted on microscope slides and fluorescence images acquired at 100x magnification.

**Statistics:** Normal distribution tests were carried out in all initial set of experiments and a non-parametric Wilcoxon test was applied accordingly. For samples with normal distribution, parametric tests include one-way ANOVA followed by a Tukey HSD or t-test. A p<0.05 was considered significant. Results are shown as mean ± s.e.m. No statistical methods were used to predetermine sample sizes. Data collection and analysis were performed blinded to the conditions of the experiments only in *in vivo* experiments. There was no randomization of data collection or processing.

## Acknowledgements

Hernandez was partially funded by a fellowship from ColFuturo. We are thankful to M. Garcia, M Dominguez, and L Guinea for technical support. This work was funded by grants SAF2010-60051/SAF2013-40710-R and by CIBERNED. AMV was funded by “Proyecto de Excelencia” from Junta de Andalucía (P08-CVI-03629). We thank TyrNovo (Israel) for their kind gift of NT219 daf2 inhibitor. We also thank NovoNordisk (Denmark) for their kind gift of S961 insulin receptor inhibitor.

## LEGENDS SUPPLEMENTARY FIGURES

**S1 Figure.**
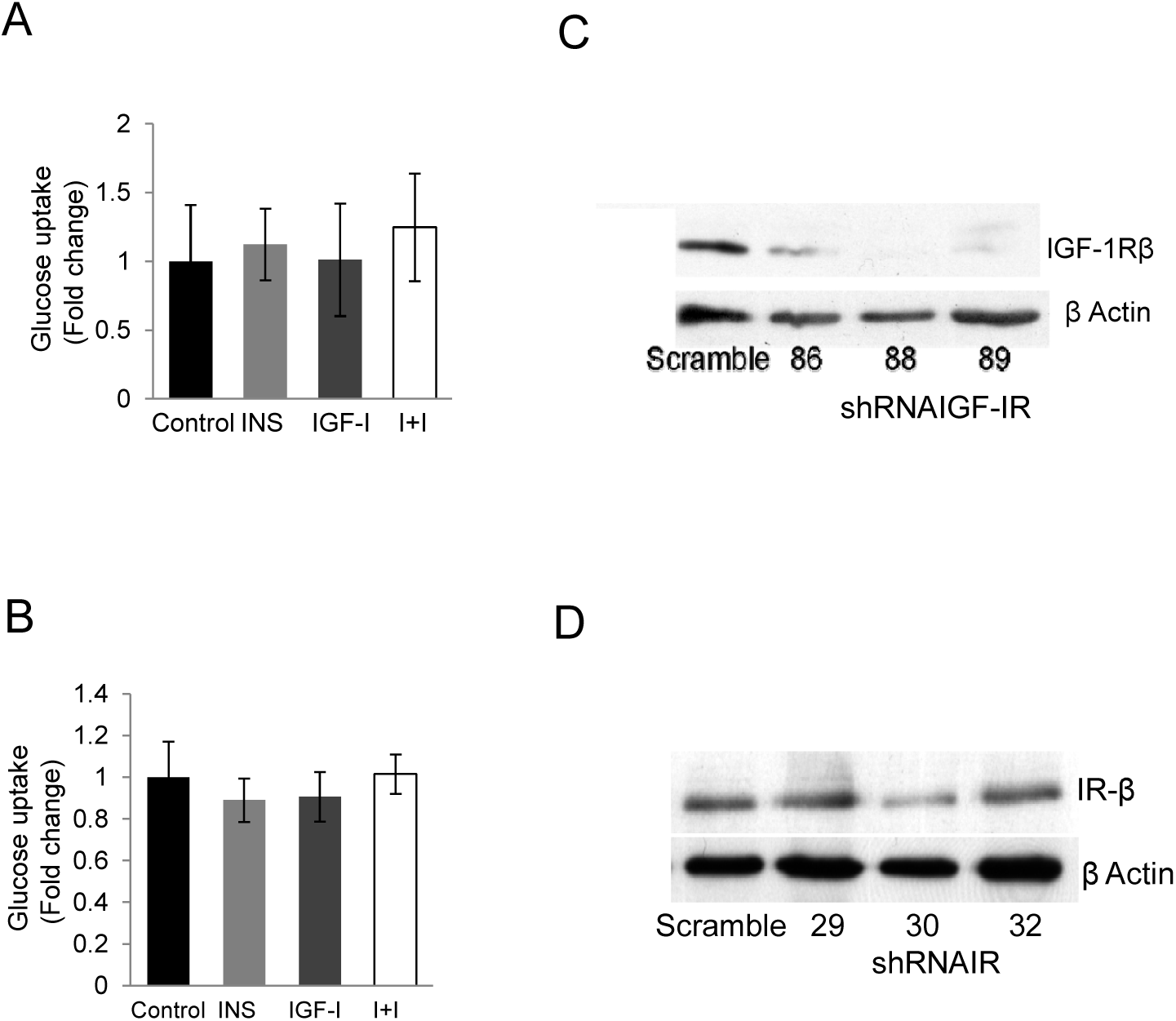
A-B, Glucose uptake in brain endothelial cells (a), and in neurons (b), determined by accumulation of the fluorescent glucose analog 6-NBDG, is not affected by co-addition of insulin and IGF-I (I+I; n=6). **C, D**, Three different shRNAs for IGF-IR (c) or for IR (d) were tested in cultured astrocytes. Number 88 for IGF-IR and number 30 for IR were selected based on their greater potency to reduce the levels of their respective targets.

**S2 Figure.**
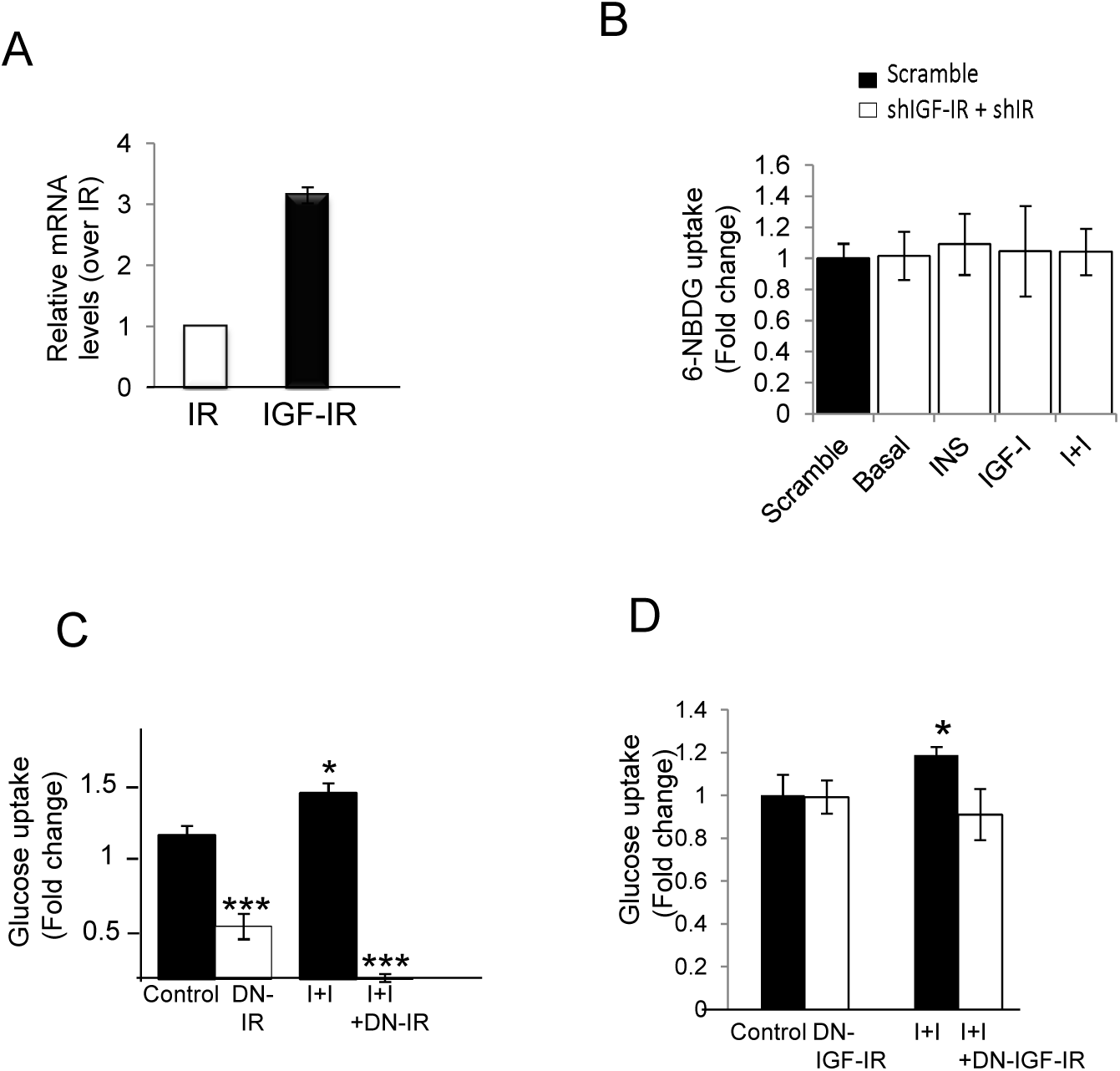
**A**, Astrocytes express both insulin (IR) and IGF-I (IGF-IR) receptors as determined by qPCR (n=6). Levels are expressed relative to IR. **B**, Interference of both IR and IGF-IR in astrocytes result in abolition of the effects of I+I and no changes in 6-NBDG uptake (n=6). **C,D**, Increased glucose uptake after I+I is blocked in astrocytes expressing either a (C) dominant negative IR, or (D) dominant negative IGF-IR (n=6 in each case; *p<0.05 and ***p<0.001 vs control).

**S3 Figure.**
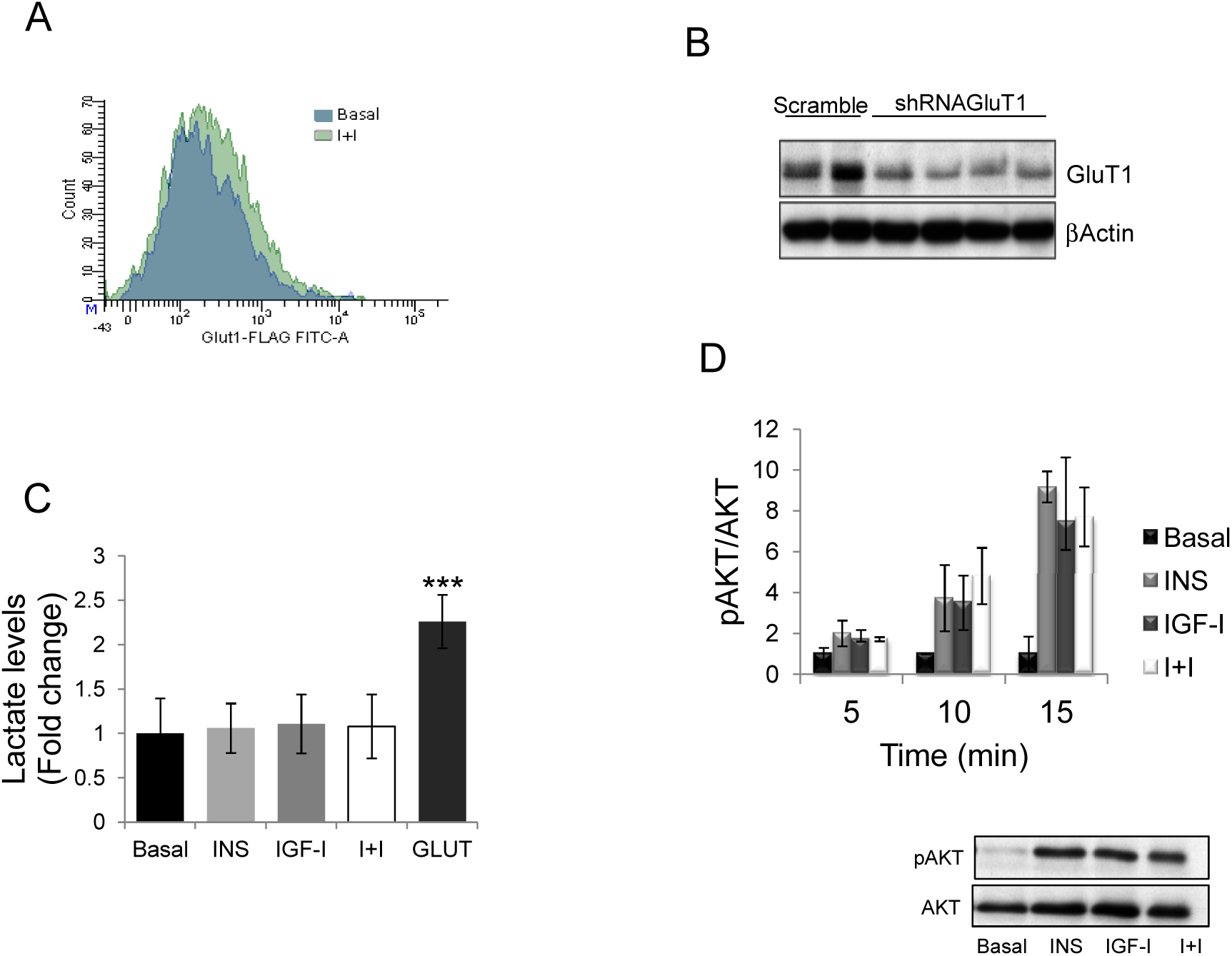
**A**, Insulin + IGF-I mobilize GLUT1 from internal compartments to the cell membrane. Representative flow cytometry chart showing an increase in GFP-tagged GLUT1^+^ astrocytes after I+I treatment as compared to basal conditions. **B**, shRNA interference of GLUT1 reduces its levels in astrocytes as compared to scramble-treated cells. **C**, Lactate levels in astrocyte cultures are not modified by insulin or IGF-I. A positive control is shown using glutamate-stimulated astrocytes (GLUT), a known stimulus of lactate production by astrocytes (n=6; ***p<0.001 vs basal). **D**, Levels of phosphorylated AKT (pAKT) were increased after either insulin or IGF-I, and co-addition of both did not elicit any further increase. Representative blot after 15 minutes of stimulation with insulin, IGF-I or both is shown. Histograms show quantification of the pAKT/AKT ratio after 5, 10, and 15 minutes of stimulation (n=3).

**S4 Figure.**
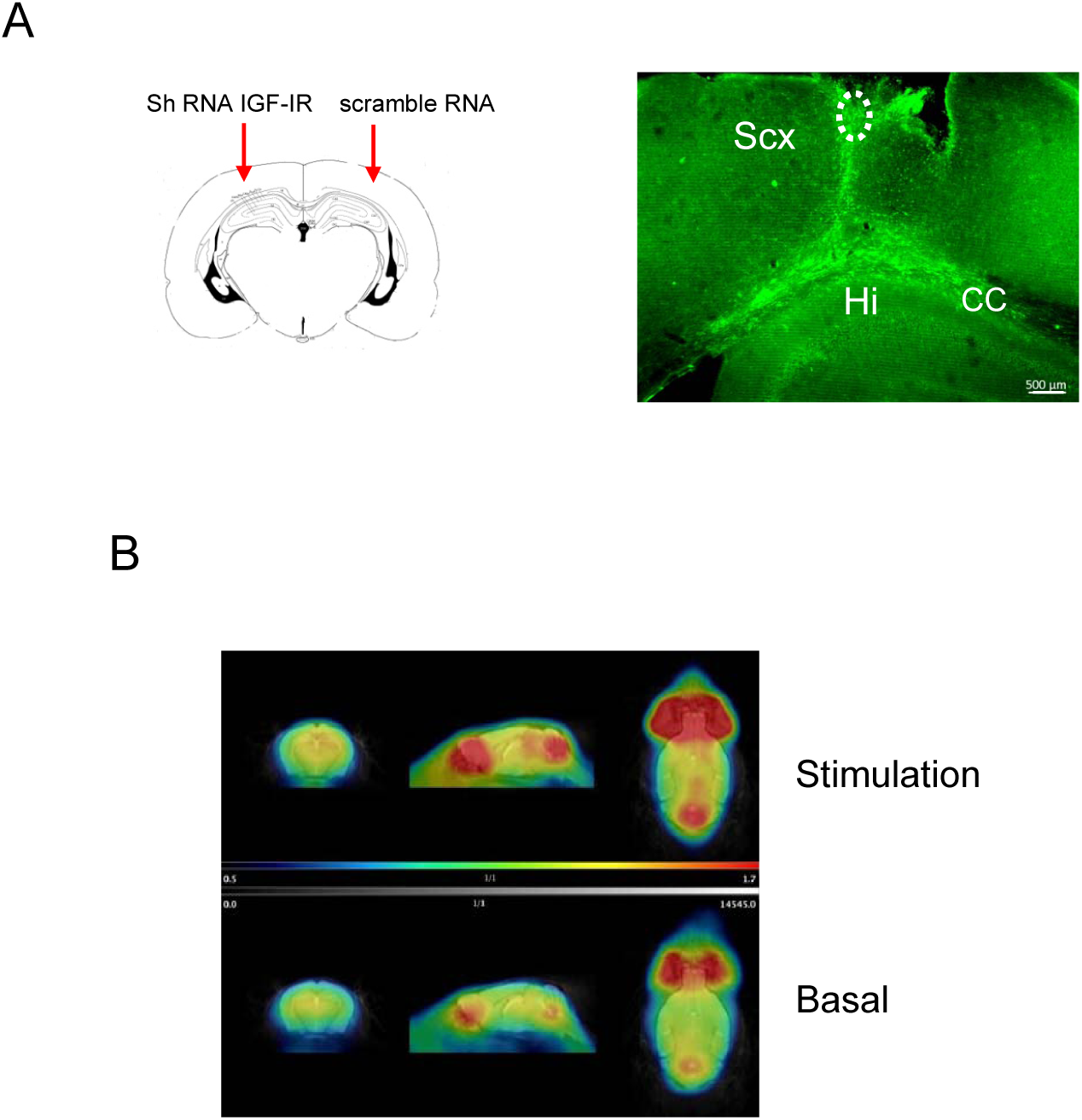
Role of IGF-IR in brain glucose uptake analyzed by PET imaging. **A**, Adult mice were injected with lentival particles expressing shRNA IGF-IR in one side of the somatosensory cortex and with scramble shRNA-expressing viral particles in the contralateral side. Right micrograph: a representative GFP staining of GFP-expressing control viral vectors is shown to illustrate the spreading of viral expression within the somatosensory cortex (Scx). The circled area depicts the region selected for microscopy analysis. Bar is 500 μm. Hi: hippocampus; CC: corpus callosum. No spreading of virus was seen in the contralateral side. **B**, Representative PET images of shRNA IGF-IR-injected mice under basal conditions and after somatosensory stimulation.

**S5 Figure.**
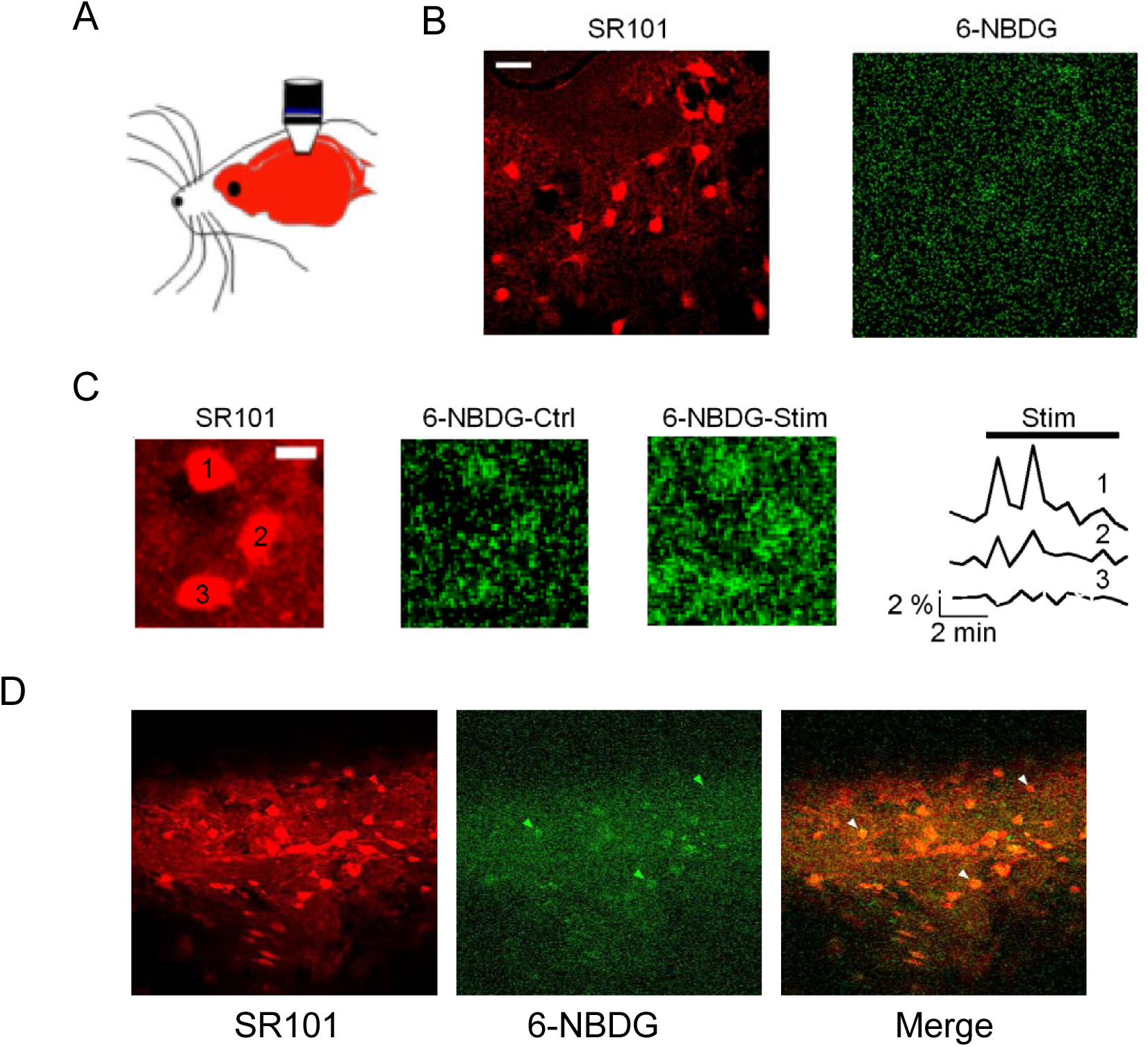
In vivo microscopy of astrocyte glucose uptake**. A**, Schematic illustration of the positioning of the microscope over the cranial window of an anesthesized mouse. **B**, Left: representative image of somatosensory cortex astrocytes labeled with SR101; right micrograph: uptake of 6NBDG in the somatosensory cortex under basal conditions. Bar is 20 μm. **C**, Representative measurements of three SR101-labeled astrocytes (1, 2, and 3 in left image) accumulating 6NBDG before (middle image), and after (right image) somatosensory stimulation. Fluorescence traces are shown in the rightmost panel. Bar is 10 μm. **D**, Representative confocal images of astrocytes (red, left image) accumulating 6NBDG (green, middle image) during whisker stimulation (see Suppl video 1 for a full temporal sequence). Three astrocytes are indicated with arrowheads, two of them accumulate more 6-NBDG as shown in the merged image (right). Astrocytes were stained with the specific astrocyte dye SR101 administered by intraperitoneal injection whereas glucose uptake was determined using the green fluorescent analog 6-NBDG delivered by intra-femoral artery injection.

**S6 Figure.**
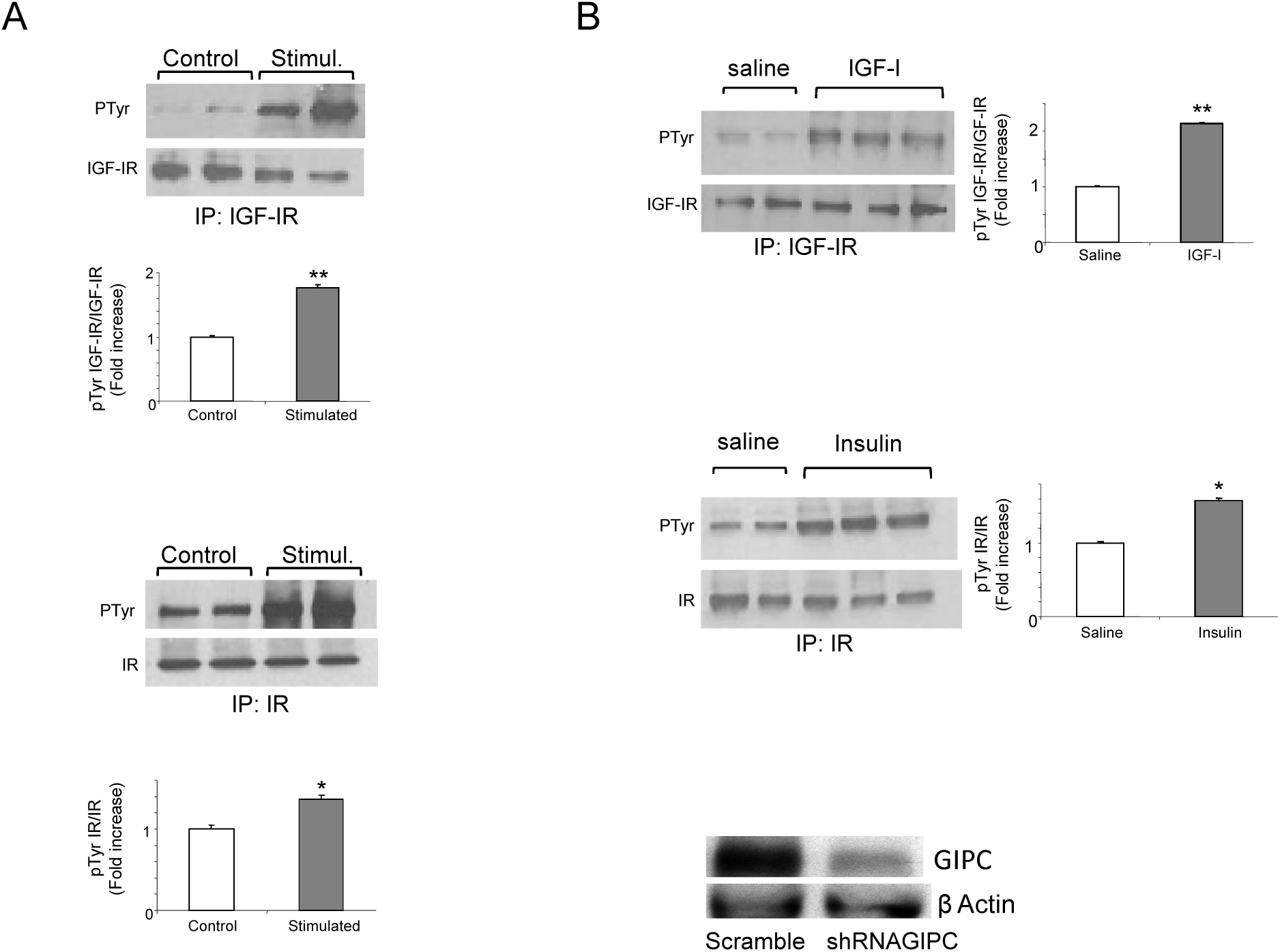
**A**, Unilateral somatosensory stimulation of the whisker pad (5Hz, 50μA during 1ms square pulse) results in enhanced phosphorylation of both the IGF-I (IGF-IR) and insulin receptors (IR) in the contralateral somatosensory cortex. Quantitation bars: **p<0.01 and *p<0.05 vs saline (n= 4). **B**, In control experiments we observed that systemic injection of 30 μg/ml of IGF-I (upper blot) or 5 U/ml insulin (lower blot) results in a similar stimulation of their respective receptors in the somatosensory cortex. Activation of the receptors is shown as amount of phosphorylated receptor normalized by total levels of immunoprecipitated corresponding receptor. Representative blots of 2 saline and 3 insulin or IGF-I-injected animals are shown (n=8 for each group).

**S7 Figure.**
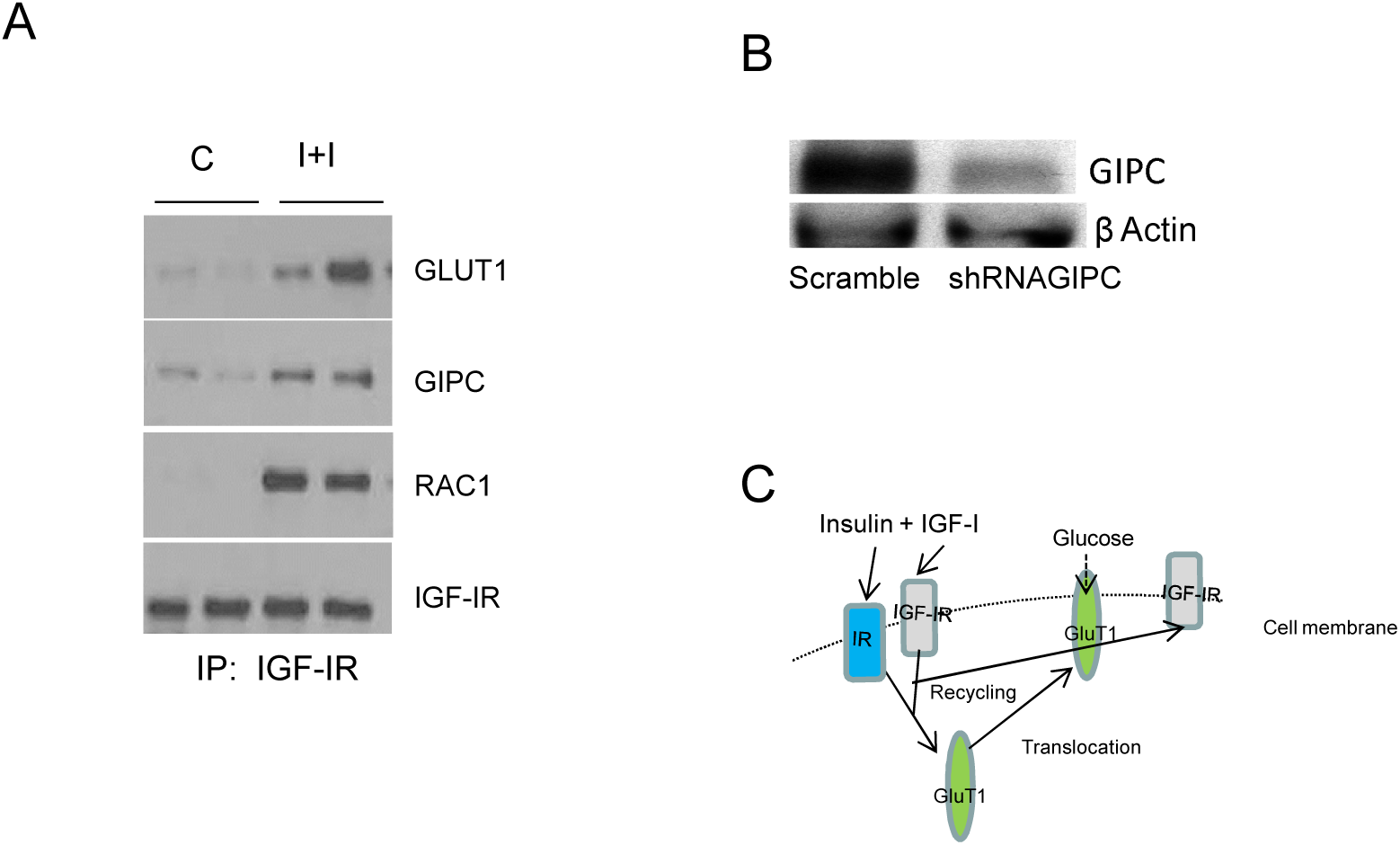
**A**, Addition of I+I increases the interaction of GLUT1 and GIPC with IGF-IR and promotes the association of Rac1 with these proteins. Co-immunoprecipitation was performed with anti-IGF-IR. **B**, shRNA GIPC reduces protein levels of GIPC in transfected astrocyte cultures. **C**, Scheme of proposed mode of action of insulin and IGF-I includes recycling of IGF-IR and translocation of GLUT1 to the cell membrane from internal compartments.

